# Efficient and Tidy Manipulation of Annotated Matrix Data with plyxp

**DOI:** 10.64898/2026.05.06.721669

**Authors:** Justin T. Landis, Michael I. Love

## Abstract

Manipulating high-dimensional omics data, such as bulk or single cell gene expression counts matrices, typically requires a bioinformatics analyst to learn domain-specific functions and syntax. These matrix-centric functions and syntax can be less intuitive than working with tidy data analytic principles, as exemplified by tools such as dplyr applied to tabular data. We propose an expressive grammar for manipulating annotated matrix data, with syntax to access, modify, and append matrix data and tabular row and column metadata, including row-wise or columnwise grouped operations. This grammar defines multiple contexts, and providing pronouns for specific recall and assignment within and across these contexts. The plyxp package is an implementation of this grammar for the R/Bioconductor ecosystem, with efficient abstractions for the SummarizedExperiment class. We demonstrate plyxp’s efficiency compared to alternative approaches on data manipulation tasks requiring computation across contexts.

## Introduction

High-throughput omics data are often summarized into one or more matrices representing counts or relative abundance of molecules in a set of experiments, as summarized across various genomic features (rows), and across different experiments or samples (columns). During subsequent analysis, it is critical to preserve links between rows and columns of the matrix data with relevant feature and sample metadata tables. For example, one may wish to divide a matrix of RNA-seq transcript counts by the transcript length (metadata associated with the transcript features along the rows) or by the library size (metadata associated with samples along the columns). This need for robust and efficient handling of annotated matrix data has motivated the development of various data classes, including but not limited to the *ExpressionSet* (Gentleman et al., 2004) *and the Summarized-Experiment* (Huber et al., 2015) in R/Bioconductor, and *AnnData* (Virshup et al., 2024) in Python. Such annotated matrix data classes and associated methods ensure that structural consistency is maintained be-tween data and metadata during common operations such as subsetting, re-ordering, and row or column binding.

Operating on these data objects requires learning domain-specific functions and syntax. Performing complex operations such as grouping over sets of rows and columns and computing further data summarization may require writing additional control structures such as for loops or using less well-known functional programming helpers, which can obfuscate the purpose of analysis code. The *tidyomics* project was created within R/Bioconductor to help address this issue (Hutchison et al., 2024), providing methods and coding paradigms for working with common R data classes representing genomic ranges (Lee et al., 2019, 2020), count matrices (Mangiola et al., 2021b), single-cell datasets (Mangiola et al., 2021a), and more. The project enables intuitive manipulation and visualization without requiring users to learn domain-specific syntax. *tidyomics* packages and workflows enable chained data-manipulative operations that resemble the style and framework of tidy data analysis (Wickham, 2014) as implemented in *dplyr* (Wickham et al., 2023) and other *tidyverse* packages, or chained operations for python *DataFrames* using *pandas* (McKinney, 2022). Taking advantage of the *tidyverse* style, *tidyomics* increases the accessibility of omics data analysis among newer R users.

Here, we propose a new expressive grammar for access and manipulation of annotated matrix data and a framework for efficiently performing the operations implied by the syntax. Data within the annotated matrix object are lazily bound to a series of environments, which means that expressions are evaluated when the user forces their symbols. Thus, a user has more freedom in how they choose to work with their data. We introduce the *plyxp* R/Bioconductor package implementing this grammar, for using *dplyr* syntax with the Bioconductor *SummarizedExperiment* (SE) data class.

The *tidySummarizedExperiment* package, which was released with Bioconductor 3.12 in 2020, also provides *dplyr* -like access to SE objects within the *tidyomics* project. The *tidySummarizedExperiment* package allows users to approach the SE as an abstracted data frame, conceptually unwrapping the compact matrix representation into a tidy, long-form version. *tidySummarizedExperiment* facilitates data exploration and visualization, allowing datasets to be directly piped into *ggplot2* plotting functions. However some complex computations involving simultaneous manipulation of matrix values and annotation metadata were slower than necessary, a limitation addressable through data masking strategies. *plyxp*’s use of data masking from the *rlang* (Henry and Wickham, 2026) *package provides gains in efficiency and is currently used to speed up existing functionality within tidySummarizedExperiment*. Further, *plyxp* reinforces the structure of the annotated matrix through additional syntax compared to *tidySummarizedExperiment*, often resulting in more verbose, but more specific evaluation.

## Syntax

We describe a grammar in which users can access, modify, and append data across different contexts of a dataset. A “context” refers to a collection of objects of the same length (for vectors) or dimension (for matrices), which collectively describe either features, samples, or a matrix of values over features and samples. For annotated matrix data with *n* rows and *p* columns, we define three contexts: the `assays` consist of one or more matrices of size *n* × *p*; the `rows` consist of row metadata (a collection of vectors of length *n* describing the features); and the cols consist of column metadata (a collection of vectors of length *p* describing the samples/cells). For example, an object representing an RNA-seq experiment might have an assays context consisting of a counts matrix and a transcripts-per-million (TPM) matrix across *n* genes and *p* samples, a rows context with feature metadata such as alternative gene identifiers and gene length, and a `cols` context with sample metadata such as library size and sample condition group.

By default, expressions are evaluated against the named symbols within the `assays` context. The user can specify to evaluate against named symbols in the row or column metadata by decorating their expressions with sentinel functions, `rows`(…) or `cols`(…), as shown in Figure 1A. Beyond communicating where expressions are evaluated, this syntax also enforces where the resulting vector or matrix will be stored. For example, with the `mutate()` command, the new objects are bound to their respective context within the object, but for the `filter()` command, instead of adding data, the syntax determines how the rows or columns are subset.

**Figure 1.**
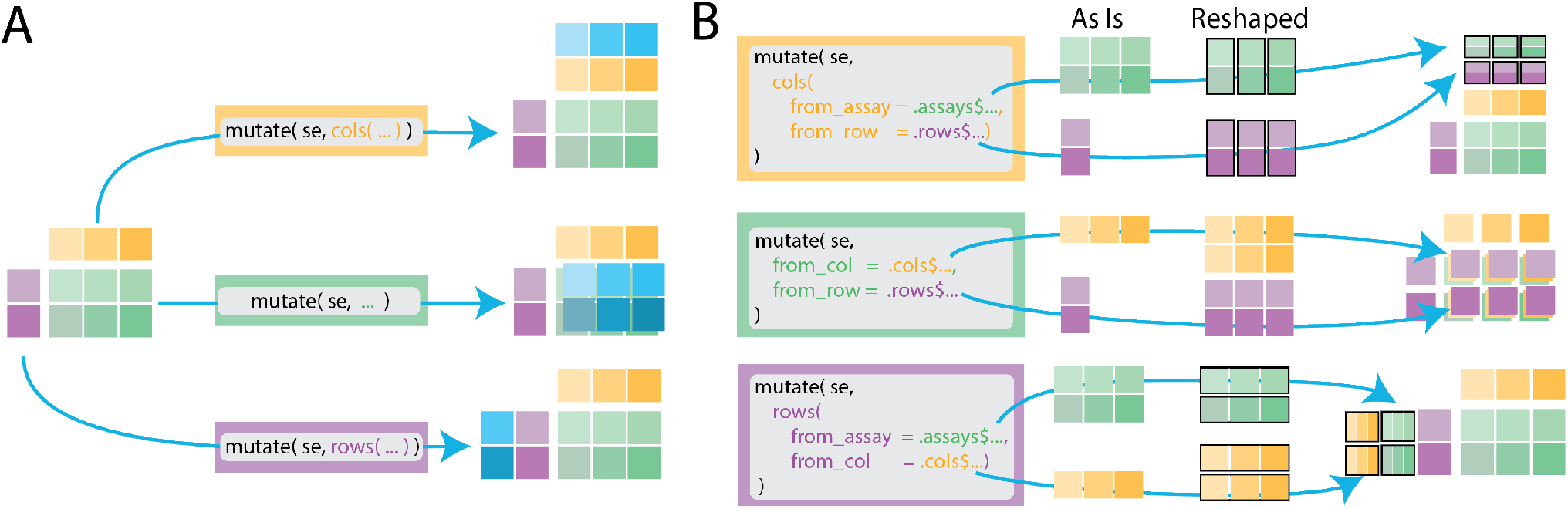
Diagram of the *plyxp* syntax with *dplyr* `mutate()` function where the `assays, rows`, and `cols` are represented by the green, purple, and yellow blocks respectively. (A) Syntax within the `mutate`(…) call determines in which contexts to insert new values (blue blocks). (B) Usage of pronouns allows multiple types of referencing between contexts. Pronouns first access the requested data “as is” before reshaping the data for the appropriate context. Black bounding boxes indicate elements of a list.

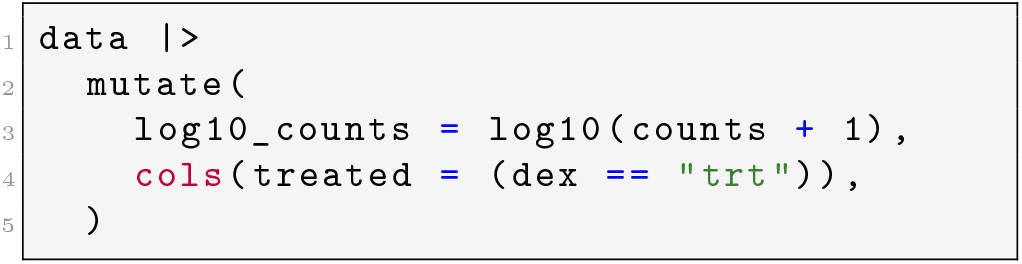

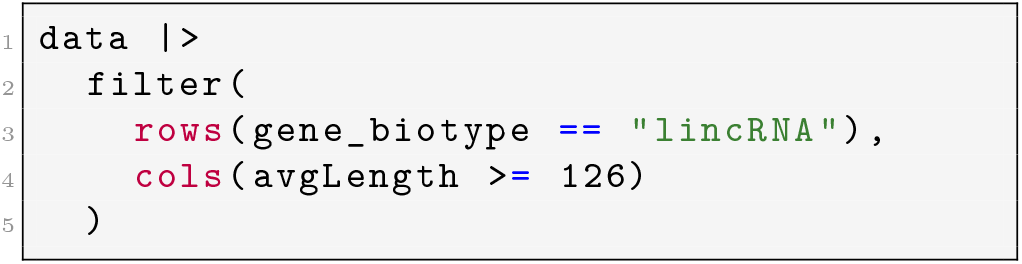

The above examples demonstrate this grammar as implemented in *plyxp*. In the first code chunk, the user creates a new matrix, the logarithm of the counts matrix plus a pseudocount, and creates a new logical variable for the column metadata, each of which are stored in their respective contexts. In the next code chunk, rows and columns are filtered based on available metadata about the genes and the samples. Here the base R |> symbol refers to piping the results of a function into the first place argument of the following function. The *plyxp* syntax is designed to be convenient and flexible, providing users with multiple tools to access or manipulate their data across contexts. For example, consider the following:

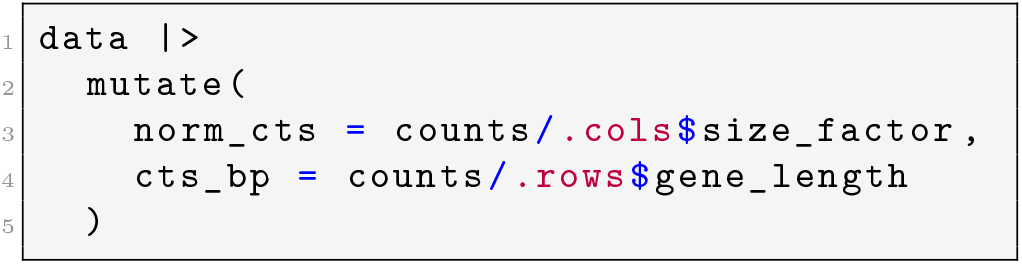

Using pronouns `.cols` and `.rows`, the user accesses vectors from another context. A “pronoun” is a special object of the *rlang* package that provides an unambiguous symbol lookup into a named list or environment (Figure 1B). In the above expression, we divide the counts matrix by vectors from the column or row metadata, replicating the column or row data to the appropriate size automatically.

Next, we consider two approaches to adding the row-wise mean of the counts matrix to the row metadata:

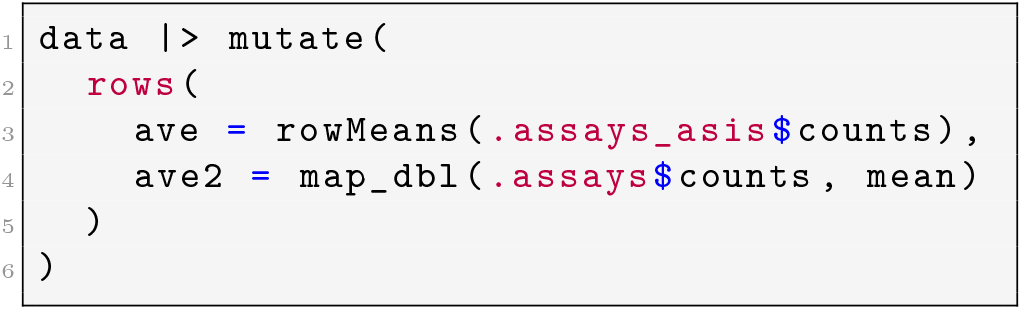

In the computation of `ave`, the user employs the efficient row-wise means operator `rowMeans`. Alternatively, for `ave2`, the expression `.assays$counts` within the row context is resolved to a list where each element is a row of the counts matrix. A list output for the `.assays` pronoun offers flexibility as lists can be supplied to the apply family of functions or using *purrr* (Wickham and Henry, 2025). In the above examples, the `dat`a object can be instantiated from any *SummarizedExperiment* using the `new_plyxp()` function (see Supplementary Note for more details).

### Semantics

*plyxp* is an implementation of this grammar for annotated matrix data stored in R/Bioconductor’s *SummarizedExperiment* class, allowing the use of *dplyr* verbs with the following expected outcomes and behaviors. *plyxp* enforces conventions that eliminate ambiguity while providing flexibility in terms of data shape and computation:

1. Expressions are scoped explicitly to an evaluation context.
2. Evaluation contexts are connected via pronouns, which provide both an `asis` and reshaped size-stable version of the requested data.
3. *dplyr* verbs (`mutate(), filter()`, etc.) only support expressions that would output the same class of object as input (SE object).

Evaluation contexts and their connection via pronouns are diagrammed in Figure 2. We note that *plyxp* relies on Bioconductor-native calls, *e*.*g*. subsetting with square brackets [,], and is thus not guaranteed to be faster than using the SE interface directly. There will always be some overhead in generating the necessary abstraction to enable implicit references into the object as required by *dplyr*. However, we aim that *plyxp* always be as close to Bioconductor-native speed as possible, with performance comparison in a following section.

**Figure 2.**
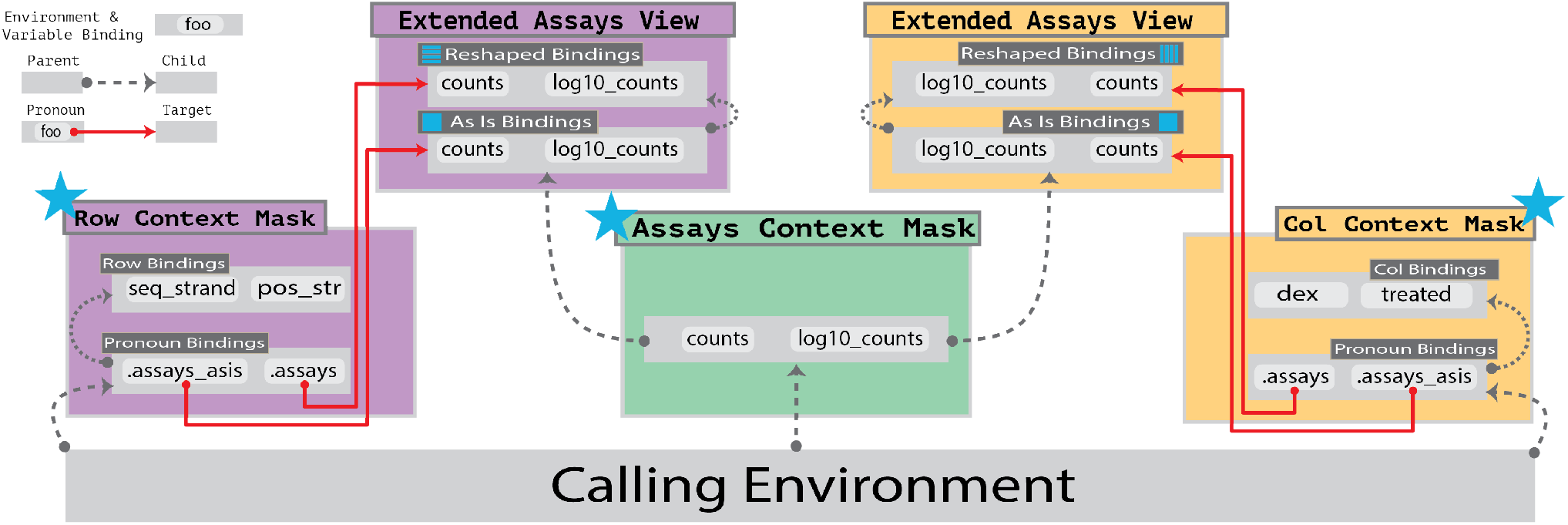
Diagram of the data masking structure in *plyxp*. Data mask contexts are grouped by color where the `assays, rows`, and `cols` contexts are represented by the green, purple, and yellow blocks respectively. Grey rectangles represent environments and contain example variable bindings. Parent environments are linked to child environments by dashed lines. Pronoun bindings are linked to their target environment contexts by red lines. For brevity, neither full environment hierarchies nor all extended environments are shown. Entry points for each context is indicated by the blue star. Through *rlang*’s tidy evaluation engine, symbols within the data mask are evaluated before searching the calling environment.

### Pronouns and Data Masks

*plyxp* enables expressive evaluation by leveraging *rlang* data masks, a series of basic R environments that are evaluated with the *rlang* tidy evaluation engine (Henry and Wickham, 2026; Wickham, 2019). *plyxp* exploits several utilities from the *rlang* package, such as lazy evaluation and *rlang* pronouns, to manage these environments. Here, “lazy evaluation” refers to expressions whose computation is delayed until the value is accessed. In *plyxp*, various expressions are lazily bound within the data mask contexts, such as the asis and reshaped bindings displayed in the extended environments of Figure 2. “*rlang* pronouns” can be conceptually thought of as a pointer to another environment. *rlang* pronouns are utilized within the *dplyr* package to disambiguate where to find a symbol, in either the calling environment or user’s data with `.env` and `.data` pronouns respectively.

In *plyxp*, these pronouns are used to connect each context through two pronouns: an asis variant and a reshaped variant. The purpose of the asis variant is to provide the user with access to the actual data stored on the object, as there may be efficient ways to operate on its native shape *e*.*g*. the row-wise and column-wise matrix operations in base R and *matrixStats* (Bengtsson, 2025). The reshaped variant transforms the requested data such that the output matches the size of the current context. The reshaped variant is intended to reduce friction from pulling data from other contexts, as there is no guarantee the target data asis will match the expected output size of the evaluation context.

With annotated matrix data, there are predictable transformations that may be used to guarantee size stability. For example, consider requesting row or column data from within the `assays` evaluation context. To make the row data size stable, we need only to replicate the requested vector *p* times in our matrix context, and then concatenate these vectors. For column data within the `assays` evaluation context, we replicate each element of the requested vector *n* times in our matrix context. Requesting assay data from within the `rows` or `cols` evaluation context, we may reshape the matrix by returning a list where each element is a row or column of the matrix, respectively. These transformations from `asis` data to the reshaped variant are diagrammed within Figure 1B.

### Grouping

In *dplyr*, specifying groups with the addition of a group_by() command listing one or more variables augments the behavior of further *dplyr* verbs, enabling computations within `mutate(), summarize(), slice()` and other commands that are iterated over data groups. *dplyr* grouping operations traditionally include additional metadata on the target object describing how the data should be partitioned. This strategy works well for data structures that are tabular, but may lead to ambiguities for data structures that deviate from this structure, such as annotated matrix data. For tabular data, there is only one dimension (along the rows) in which those data can be partitioned, and no matter how the rows are partitioned, a new table can always be constructed from the results. This is not true for matrices; consider an attempt to group by matrix values > 100. Here, there is no guarantee the partitioning of arbitrary elements of a matrix would create valid sub-matrices. It is for this reason that *plyxp* follows strict heuristics for grouping and only allows groupings that partition rows or columns of the matrix.

This restriction coincides with the natural extension of applying grouping to tabular data from *dplyr*. We may conceptually regard the row data and the column data as independent and grouped as normal tabular structures. The row data and the column data are related to one another through the matrix, and thus the matrix observes all grouping partitions imposed on both dimensions. Since the row data and the column data groupings are independent, the `rows` and `cols` contexts only consider their own respective grouping during evaluation. In other words, the `assays` context will always respect groupings about the `rows` and `cols` contexts, but the rows context drops any cols grouping and `cols` context drops any `rows` grouping.

### Performance Comparison

All evaluation within *plyxp* is intended to take place within one of the specified contexts mentioned previously. Due to the particular shape of annotated matrix data, each context has a unique notion of size and a unique set of symbols within. Additional syntax is used to declare where an expression evaluates and there is no additional cost to accessing data asis. An alternative to this approach would be to force all variables within the same context, such as an instantiated or abstracted table where all contexts are joined together. A single context would require variables within the assays to be unwound into one-dimensional vectors, and data within the row and column metadata tables to be replicated appropriately.

We compared the performance of various approaches to the operation of scaling the counts matrix by gene length using the *airway* dataset (Himes et al., 2014). For this operation, *plyxp* was faster than the unwinding approach but somewhat slower than native calls (Table 1). We also compared computing row means with either the asis or reshaped pronouns using either the optimized `rowMeans()` or `map_dbl()` from *purrr*, respectively. Naturally, opting to not reshape the data yielded substantial computation time saving. Evaluation code available in the comparison.R file within the associated GitHub repository (Landis and Love, 2026). Although *plyxp* introduces slight overhead, we find that its gains in usability and syntactic precision justify the cost. Furthermore, for more compute-heavy operations, especially in grouping scenarios, *plyxp* performs at least as well as native calls. Additional performance considerations are discussed in more detail in the Supplemental Note.

**Table 1.**
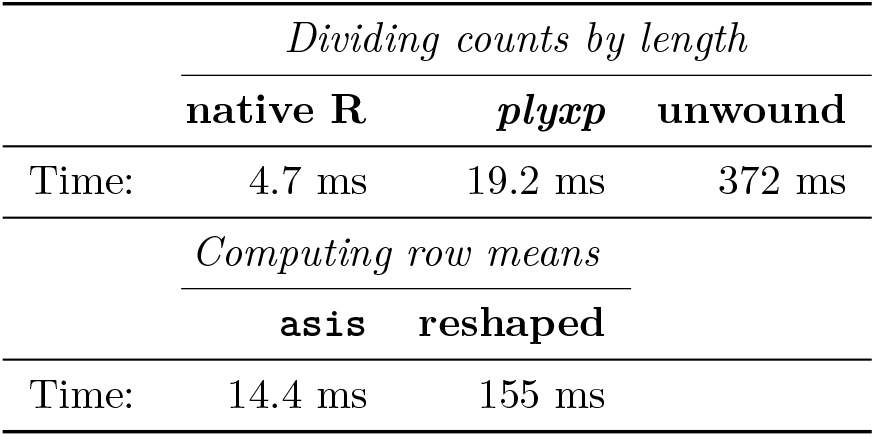
Performance comparison of *plyxp* and alternative approaches. Run times reported as median over repeated evaluations. Note that the “Computing row means” task compares pronoun access options within *plyxp* as demonstrated in the last code chunk of the Syntax section.

## Conclusion

*plyxp* provides a framework for operating on annotated matrix data, making common *dplyr* functions as natural to use as with tabular data. Using data masking from the *rlang* package, we introduce novel syntax for operating on SE objects within Bioconductor that is ergonomically inspired by the *dplyr* style and is still unambiguously expressive. *tidySummarizedExperiment* has incorporated *plyxp* methods internally for some operations as of Bioconductor 3.22 release in Fall 2025, such that users of both *plyxp* and *tidySummarizedExperiment* benefit from the performance gains described above, as compared to operations that would require creation of a long form tabular representation of the annotated matrix data. We imagine that the grammar described here could be applied to manipulate *AnnData* with *pandas* methods. In the future, we plan to release the internals of *plyxp* that construct and manage the data masks so that developers can choose to create their own *dplyr* style for additional R classes.

## Supporting information

Supplemental plyxp Usage Vignette

## Code Availability

The *plyxp* R package is available as part of Bioconductor since version 3.20, installable with BiocManager::install(“plyxp”), and at https://github.com/jtlandis/plyxp.

All code used in this paper, including comparisons of *plyxp* to alternative implementations, and reproducible code chunks appearing in this manuscript, is available at https://github.com/jtlandis/plyxp_paper (Landis and Love, 2026). Evaluations were run with Bioconductor 3.23 and R 4.6.0, with the following package versions: *plyxp* (1.6.1), *dplyr* (1.2.1), *tidySummarizedExperiment* (1.22.0), *airway* (1.32.0).

## Author contributions statement

J.T.L. developed the software. J.T.L. and M.I.L. contributed to the conceptualization and design of the software and to the writing of the manuscript.

## Acknowledgments

The authors thank Rachel Sharp, Beatriz Campillo Miñano, Stuart Lee, and Stefano Mangiola for comments on an earlier draft.

## Supplementary Note

### Interoperability with *tidySummarizedExperiment*

*plyxp* enables users to continue using the *tidySummarizedExperiment* package for operating directly on SE objects using *dplyr* verbs without namespace conflicts, as it employs a class *PlySummarizedExperiment* to trigger its methods. In practice, the user instantiates a *PlySummarizedExperiment* object by providing an SE to the `new_plyxp()` function. The *PlySummarizedExperiment* class is related to the SE class by composition in that *PlySummarizedExperiment* contains a single slot which accepts any object inheriting from the *SummarizedExperiment* class. Due to composition, *plyxp* can generalize its core functionalities to more specialized sub classes of the *SummarizedExperiment* class, such as *SingleCellExperiment* class. Meanwhile, *tidySummarizedExperiment* users can continue to use *dplyr* verbs as previously without any change in behavior.

### Performance Considerations

Abstracting a user’s entry point into annotated matrix objects provides several advantages. To emphasize these advantages, we will first consider a counterexample. We will require an isomorphic implementation, meaning that both input and output are the same object classes. A sensible approach is to unwind the annotated matrix into a tabular structure and then reconstruct the original annotated matrix before returning. An advantage of this framework is that it allows the use of standard tabular processing libraries like *dplyr* or *pandas* directly. However, it has notable drawbacks that depend on the complexity of the user’s object.

Under this unwinding model, one can expect a large amount of overhead dedicated to transforming the object into the tabular structure required for a tabular processing library. In the worst case scenario, this transformation is done eagerly for all data in the object per method, which would be a significant amount of work done before any user expressions are evaluated. Another consideration for constructing this tabular structure would be the possibility of symbol collision, that is, how to resolve the resulting column names if the input contexts have identical names. In order to work, some heuristic would need to be established such as a renaming convention or context precedence. In either case in which these collisions would occur, the resulting code would be ambiguous, and it may be better to send an error and enforce uniqueness across the entire object than force the user to handle heuristic name resolution. The final issue with the conversion is the loss of information regarding where data is from and where it should be set. In the case of *dplyr* or *pandas* verbs that modify the object, `mutate()` or `.assign()` respectively, it is unclear where new data should be set in the reconstructed annotated matrix object. A safe choice would be to set the new data as a new assay, however, if the implementation would need to set the result in either the row or column metadata tables, then additional computation would be required to ensure the resulting new vector’s uniqueness and its ability to fit in one of those positions.

In *plyxp*, all issues related to ambiguity are resolved by constructing three separate data masks. *plyxp* imposes a verbose, but specific syntax to communicate intent of which context an expression should be evaluated. Under this model, there is no additional computational cost for direct access, no possibility of symbol collision, and no ambiguity about where data will be set on the output object. Only symbols bound within the contexts the user requests are evaluated, saving time compared to approaches that manipulate the entire structure all together. If the user requires cross-context data in which the requested data may be transformed, these bindings are also lazy and save computation time if never forced. This is especially convenient for grouping operations, in which constructing groups eagerly for a dense object would require extensive computation. In this model’s design of the data masks, the cost of performing this computation is only paid once per data mask per symbol.

Finally, *plyxp* makes some assumptions about how the user may want their data to be shaped in a given context, but provides the user with a way to access the underlying data without reshaping. This is motivated by the fact that, despite utilizing efficient subset functions when possible, it may still be more efficient for the user to operate on the native data structure with optimized routines, opting to forego any potentially expensive transformations.

