## Supplemental plyxp Usage Vignette for "Efficient and Tidy Manipulation of Annotated Matrix Data with plyxp"

Justin Landis Michael Love

2026-05-06

### Table of contents

1 Introduction 1

2 Load an RNA-seq data object 2

3 Basic manipulation of RNA-seq data 2

4 Exploring data alongside DE results 5

5 Computing and plotting abundance 7

6 Grouping and summarizing abundance over samples 8

7 Session Info 12

### 1 Introduction

This document is a modified version of the *plyxp* vignette as a supplement for the *plyxp* paper. The `.qmd` file that generated this document can be found here:

https://github.com/jtlandis/plyxp_paper

The full and up-to-date *plyxp* vignette can be found here:

https://jtlandis.github.io/plyxp/articles/plyxp.html

To install *plyxp* use the following code:

```
if (!require("BiocManager", quietly = TRUE))
    install.packages("BiocManager")
BiocManager::install("plyxp")
```

Information about *plyxp* can be found at the following two locations:

- https://github.com/jtlandis/plyxp
- https://bioconductor.org/packages/plyxp

### 2 Load an RNA-seq data object

We load the *airway* dataset which contains a *SummarizedExperiment* object with gene-level counts across eight samples. The *airway* dataset summarizes an RNA-Seq experiment on four human airway smooth muscle cell lines, both as controls and treated with dexamethasone. Details on the gene model and read counting procedure are provided in the *airway* Bioconductor package vignette. For more details on construction of *SummarizedExperiment* (SE) objects, see the vignette for the package of the same name, or the *tximeta* package which can construct these objects from output of common RNA-seq quantification tools.

The citation for the experiment is:

> Himes BE, Jiang X, Wagner P, Hu R, Wang Q, Klanderman B, Whitaker RM, Duan Q, Lasky-Su J, Nikolos C, Jester W, Johnson M, Panettieri R Jr, Tantisira KG, Weiss ST, Lu Q. "RNA-Seq Transcriptome Profiling Identifies CRISPLD2 as a Glucocorticoid Responsive Gene that Modulates Cytokine Function in Airway Smooth Muscle Cells." *PLOS One*. 2014 Jun 13;9(6):e99625. PMID: 24926665. GEO: GSE52778.

```
library(airway)
data(airway)
```

First we prepare the `airway` object a bit, by removing extra variables from the row and column metadata.

```
# cleaning up the metadata...
colData(airway) <- colData(airway)[,c("cell","dex")]
rowData(airway) <- rowData(airway)[,c("gene_id","symbol","gene_biotype")]
rowData(airway)$gene_length <- sum(width(reduce(rowRanges(airway))))
rowRanges(airway) <- NULL # ranges not needed for this demo, simplifying...
```

### 3 Basic manipulation of RNA-seq data

We initialize the dataset for use with *plyxp* by calling `new_plyxp()` on an SE object. This creates a *PlySummarizedExperiment*, a simple wrapper class that contains the SE, and changes the behavior of the dplyr verbs when they are applied to our new object.

```
library(plyxp)
xp <- new_plyxp(airway)
class(xp)
```

```
[1] "PlySummarizedExperiment"
attr(,"package")
[1] "plyxp"
```

We can perform one or more operations, modifying the contents of the object:

```
xp |>
  mutate(
    log10_cts = log10(counts + 1),
    cols(treated = dex == "trt"),
    rows(gene_kb = gene_length / 1e3)
  )
```

```
# A SummarizedExperiment-tibble Abstraction: 63,677 × 8
    .features       .samples   | counts log10_cts | gene_id  symbol gene_biotype
    <chr>           <chr>      |  <int>     <dbl> | <chr>    <chr>  <chr>
  1 ENSG00000000003 SRR1039508 |    679      2.83 | ENSG000… TSPAN6 protein_cod…
  2 ENSG00000000005 SRR1039508 |      0      0    | ENSG000… TNMD   protein_cod…
  3 ENSG00000000419 SRR1039508 |    467      2.67 | ENSG000… DPM1   protein_cod…
  4 ENSG00000000457 SRR1039508 |    260      2.42 | ENSG000… SCYL3  protein_cod…
  5 ENSG00000000460 SRR1039508 |     60      1.79 | ENSG000… C1orf… protein_cod…
  …         …             …          …         …        …       …         …
n-4 ENSG00000273489 SRR1039521 |      0      0    | ENSG000… RP11-… antisense
n-3 ENSG00000273490 SRR1039521 |      0      0    | ENSG000… TSEN34 protein_cod…
n-2 ENSG00000273491 SRR1039521 |      0      0    | ENSG000… RP11-… lincRNA
n-1 ENSG00000273492 SRR1039521 |      0      0    | ENSG000… AP000… lincRNA
n   ENSG00000273493 SRR1039521 |      0      0    | ENSG000… RP11-… lincRNA
#   n = 509,416
#   6 more variables: gene_length <int>, gene_kb <dbl>, `` <>, cell <fct>,
#   dex <fct>, treated <lgl>
```

The above computations only made use of data *within* each context. Using pronouns, we can also access data *across* contexts:

```
xp |>
  mutate(
    cols(col_sum_per_mil = colSums(.assays_asis$counts)/1e6),
    scaled_counts = counts/.cols$col_sum_per_mil,
    rows(ave_scl_cts = rowMeans(.assays_asis$scaled_counts))
  )
```

```
# A SummarizedExperiment-tibble Abstraction: 63,677 × 8
    .features      .samples | counts scaled_counts | gene_id symbol gene_biotype
    <chr>          <chr>    |  <int>         <dbl> | <chr>   <chr>  <chr>
  1 ENSG000000000… SRR1039… |    679         32.9  | ENSG00… TSPAN6 protein_cod…
  2 ENSG000000000… SRR1039… |      0          0    | ENSG00… TNMD   protein_cod…
  3 ENSG000000004… SRR1039… |    467         22.6  | ENSG00… DPM1   protein_cod…
  4 ENSG000000004… SRR1039… |    260         12.6  | ENSG00… SCYL3  protein_cod…
  5 ENSG000000004… SRR1039… |     60          2.91 | ENSG00… C1orf… protein_cod…
  …         …            …          …           …        …       …         …
n-4 ENSG000002734… SRR1039… |      0          0    | ENSG00… RP11-… antisense
n-3 ENSG000002734… SRR1039… |      0          0    | ENSG00… TSEN34 protein_cod…
n-2 ENSG000002734… SRR1039… |      0          0    | ENSG00… RP11-… lincRNA
n-1 ENSG000002734… SRR1039… |      0          0    | ENSG00… AP000… lincRNA
n   ENSG000002734… SRR1039… |      0          0    | ENSG00… RP11-… lincRNA
#   n = 509,416
```

```
#   6 more variables: gene_length <int>, ave_scl_cts <dbl>, `` <>, cell <fct>,
#   dex <fct>, col_sum_per_mil <dbl>
```

We can also perform grouping operations, e.g. here grouping by the gene biotype, and computing the sum of counts for each sample:

```
summary <- xp |>
  group_by(rows(gene_biotype)) |>
  summarize(
    col_sums = colSums(counts),
    # may rename rows with .features
    rows(.features = unique(gene_biotype))
  )
```

Filtering to a single sample, and examining the first six biotypes:

```
summary |>
  filter(cols(.samples == "SRR1039508")) |>
  head()
```

```
# A SummarizedExperiment-tibble Abstraction: 6 × 1
# Groups: rows(gene_biotype)
  .features              .samples   | col_sums | gene_biotype       | cell  dex
  <chr>                  <chr>      |    <dbl> | <chr>              | <fct> <fct>
1 protein_coding         SRR1039508 | 19413626 | protein_coding     | N613… untrt
2 pseudogene             SRR1039508 |   807285 | pseudogene         | N613… untrt
3 processed_transcript   SRR1039508 |    47547 | processed_transc… | N613… untrt
4 antisense              SRR1039508 |    49682 | antisense          | N613… untrt
5 lincRNA                SRR1039508 |   133335 | lincRNA            | N613… untrt
6 polymorphic_pseudogene SRR1039508 |     2804 | polymorphic_pseu… | N613… untrt
```

We can convert the SE to tibble after selecting one of the assays (in this case there is only one assay):

```
library(tibble)
summary |>
  pull(col_sums) |>
  as_tibble(rownames = "type") |>
  head()
```

```
# A tibble: 6 × 9
  type         SRR1039508 SRR1039509 SRR1039512 SRR1039513 SRR1039516 SRR1039517
  <chr>             <dbl>      <dbl>      <dbl>      <dbl>      <dbl>      <dbl>
1 protein_cod…   19413626   17741060   23926011   14360299   23003444   29233398
2 pseudogene       807285     733916     868950     478980     852106     946776
3 processed_t…      47547      46534      47788      26367      43965      52063
4 antisense         49682      43769      60098      33066      56696      66335
5 lincRNA          133335     120060     206075     125015     145078     170641
6 polymorphic…       2804       2895       3417       2247       3497       4065
#   2 more variables: SRR1039520 <dbl>, SRR1039521 <dbl>
```

To extract the *SummarizedExperiment* from a *PlySummarizedExperiment* we just use the `se()` function:

```
se(xp)
```

```
class: SummarizedExperiment
dim: 63677 8
metadata(1): ''
assays(1): counts
rownames(63677): ENSG00000000003 ENSG00000000005 ... ENSG00000273492
  ENSG00000273493
rowData names(4): gene_id symbol gene_biotype gene_length
colnames(8): SRR1039508 SRR1039509 ... SRR1039520 SRR1039521
colData names(2): cell dex
```

### 4 Exploring data alongside DE results

Below we demonstrate running differential expression (DE) analysis on the SE object, and then manipulating the SE alongside results including p-values and log2 fold changes (LFC).

```
# define minimal expression as count >= 10
filt_xp <- xp |>
  mutate(rows(
    min_exprs_10 = rowSums(.assays_asis$counts >= 10)
  )) |>
  filter(rows(min_exprs_10 >= 4)) |>
  mutate(cols(
    dex = factor(dex, levels=c("untrt","trt"))
  ))
```

The following code passes the SE object to the *DESeq2* functions, creating a *DESeqDataSet* and then running a likelihood ratio test. This is standard R/Bioconductor code, showing that *plyxp* can easily be used alongside other Bioconductor packages.

```
library(DESeq2)
dds <- se(filt_xp) |>
  DESeqDataSet(design=~cell + dex) |>
  DESeq(test="LRT", reduced=~cell, quiet=TRUE)
```

We pick a subset of the columns of results in the `rowData` to use in further analysis:

```
cols_to_save <- c(
  "gene_id","symbol","gene_biotype","gene_length",
  "baseMean","dispersion","dex_trt_vs_untrt","LRTPvalue"
)
rowData(dds) <- rowData(dds)[,cols_to_save]
assays(dds) <- assays(dds)[c("counts","mu")]
```

We can easily pass back into *plyxp* by wrapping the *DESeqDataSet* as a *PlySummarizedExperiment*. This strategy works for any classes that descend from SE (e.g. *DESeqDataSet*, *SingleCellExperiment*, etc.).

We then compute adjusted p-values and the sign of the LFC.

```
de_xp <- dds |> new_plyxp()
de_xp <- de_xp |>
  mutate(rows(
    padj = p.adjust(LRTPvalue, method="BH"),
    lfc_sign = sign(dex_trt_vs_untrt)
  ))
rowData(de_xp) |> head()
```

```
DataFrame with 6 rows and 10 columns
                        gene_id      symbol   gene_biotype gene_length
                    <character> <character>    <character>   <integer>
ENSG00000000003 ENSG00000000003      TSPAN6 protein_coding        2968
ENSG00000000419 ENSG00000000419        DPM1 protein_coding        1207
ENSG00000000457 ENSG00000000457       SCYL3 protein_coding        6876
ENSG00000000460 ENSG00000000460    C1orf112 protein_coding        6354
ENSG00000000971 ENSG00000000971         CFH protein_coding        8144
ENSG00000001036 ENSG00000001036       FUCA2 protein_coding        3119
                 baseMean dispersion dex_trt_vs_untrt   LRTPvalue        padj
                <numeric>  <numeric>        <numeric>   <numeric>   <numeric>
ENSG00000000003  710.0340 0.00820637       -0.3852374 1.29180e-04 9.95627e-04
ENSG00000000419  521.2139 0.00999288        0.2028550 6.90960e-02 1.87545e-01
ENSG00000000457  237.6016 0.01489069        0.0339708 8.09961e-01 9.08055e-01
ENSG00000000460   58.0405 0.05701636       -0.0935336 7.39196e-01 8.69040e-01
ENSG00000000971 5827.0185 0.00744417        0.4224097 2.27453e-06 2.59976e-05
ENSG00000001036 1284.7322 0.00687809       -0.2450451 6.14104e-03 2.80279e-02
                 lfc_sign
                <numeric>
ENSG00000000003        -1
ENSG00000000419         1
ENSG00000000457         1
ENSG00000000460        -1
ENSG00000000971         1
ENSG00000001036        -1
```

Getting a sense of the number of genes at FDR $< 5\%$:

```
with(rowData(de_xp),
  table(lfc_sign, sig_5pct = padj < .05)
)
```

```
        sig_5pct
lfc_sign FALSE TRUE
      -1  6402 1872
      1   5688 2177
```

### 5 Computing and plotting abundance

In the following section, we compute abundance as *transcripts per million* (TPM). This involves scaling the observed counts by the gene length (here we have access to the number of exonic base pairs), and then dividing by the column sum to account for sequencing depth differences. The resulting matrix is multiplied by 1 million, such that the columns sum to 1 million.

```
de_tpm <- de_xp |>
  mutate(
    RPK = counts / (.rows$gene_length / 1e3),
    cols(rpk_scale = colSums(.assays_asis$RPK) / 1e6),
    TPM = RPK / .cols$rpk_scale
  )

colSums(assay(de_tpm, "TPM"))
```

```
SRR1039508 SRR1039509 SRR1039512 SRR1039513 SRR1039516 SRR1039517 SRR1039520
     1e+06      1e+06      1e+06      1e+06      1e+06      1e+06      1e+06
SRR1039521
     1e+06
```

We can directly pipe an SE object into a `ggplot()` command after loading *tidySummarizedExperiment*. We can use this to plot our newly computed TPM values across genes and samples, splitting by gene biotype, and by the treatment group.

```
library(tidySummarizedExperiment)
library(ggplot2)

biotypes <- c("protein_coding", "lincRNA", "antisense")
de_tpm |>
  filter(rows(gene_biotype %in% biotypes)) |>
  se() |>
  ggplot(aes(x = TPM + 0.1, y = gene_biotype)) +
  geom_violin(aes(fill = dex), color = NA) +
  scale_x_log10(labels = scales::label_log())
```

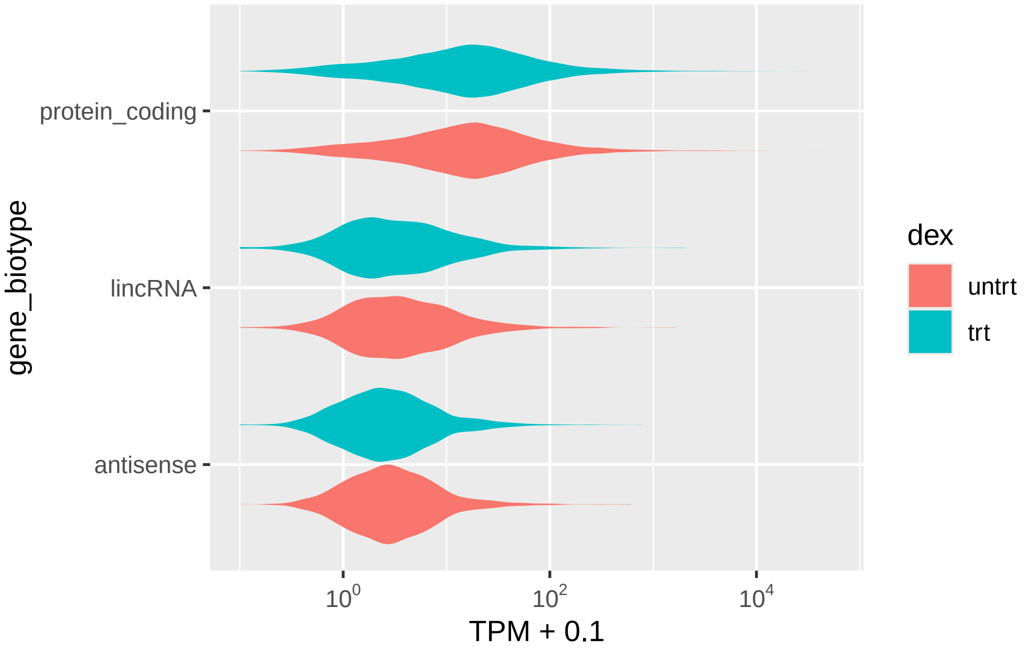


### 6 Grouping and summarizing abundance over samples

Suppose we want to make some plots of the abundance (TPM) in each treatment group, focusing on the genes with evidence of differential expression. First we filter to significant genes and group samples by condition group, then we can summarize the average TPM in each group, with the following code:

```
de_tpm_summary <- de_tpm |>
  filter(rows(padj < 0.05)) |>
  group_by(cols(dex)) |>
  summarize(ave_TPM = rowMeans(TPM), cols(.samples = unique(dex)))

de_tpm_summary
```

```
# A SummarizedExperiment-tibble Abstraction: 4,049 × 2
# Groups: cols(dex)
    .features       .samples | ave_TPM | gene_id symbol gene_biotype gene_length
    <chr>           <chr>    |   <dbl> | <chr>   <chr>  <chr>              <int>
  1 ENSG00000000003 untrt    |   46.9  | ENSG00… TSPAN6 protein_cod…        2968
  2 ENSG00000000971 untrt    |  105.   | ENSG00… CFH    protein_cod…        8144
  3 ENSG00000001036 untrt    |   77.6  | ENSG00… FUCA2  protein_cod…        3119
  4 ENSG00000001167 untrt    |   19.7  | ENSG00… NFYA   protein_cod…        3811
  5 ENSG00000001629 untrt    |   46.8  | ENSG00… ANKIB1 protein_cod…        7130
  …         …         …          …         …       …         …               …
n-4 ENSG00000272870 trt      |   11.9  | ENSG00… RP11-… antisense           2495
n-3 ENSG00000273038 trt      |    9.40 | ENSG00… RP11-… lincRNA             2051
n-2 ENSG00000273131 trt      |   21.1  | ENSG00… RP6-4… processed_t…        3827
n-1 ENSG00000273179 trt      |   39.0  | ENSG00… RP11-… antisense            397
n   ENSG00000273290 trt      |    9.40 | ENSG00… CTC-2… lincRNA             3533
```

```
#   n = 8,098
#   8 more variables: baseMean <dbl>, dispersion <dbl>, dex_trt_vs_untrt <dbl>,
#   LRTPvalue <dbl>, padj <dbl>, lfc_sign <dbl>, `` <>, dex <fct>
```

Next we group the data by gene biotype and select the top three genes according to the largest `ave_TPM` difference between treatment and control samples. This gives us back three genes for each gene biotype (except for miRNA which only has two significant genes).

This can be done in two ways: since we know there are only two treatment groups, we may use `purrr::map_dbl` on the reshaped variant of `ave_TPM`, which returns a list of the rows of the `ave_TPM` matrix. In other words, the reshaped data will be a list of vectors of length two (the average TPM for samples in each treatment group).

```
de_tpm_summary |>
  group_by(rows(gene_biotype)) |>
  arrange(rows(
    purrr::map_dbl(
      # list of length 2
      # numeric vectors
      .assays$ave_TPM,
      ~ abs(diff(.x))
    ) |>
      dplyr::desc()
  )) |>
  slice(rows(1:3))
```

```
# A SummarizedExperiment-tibble Abstraction: 26 × 2
# Groups: rows(gene_biotype)
    .features       .samples | ave_TPM | gene_biotype gene_id symbol gene_length
    <chr>           <chr>    |   <dbl> | <chr>        <chr>   <chr>        <int>
  1 ENSG00000087086 untrt    |  4.47e4 | protein_cod… ENSG00… FTL            878
  2 ENSG00000075624 untrt    |  2.76e3 | protein_cod… ENSG00… ACTB          3256
  3 ENSG00000103888 untrt    |  1.79e3 | protein_cod… ENSG00… KIAA1…        7621
  4 ENSG00000269378 untrt    |  2.26e3 | pseudogene   ENSG00… ITGB1…         609
  5 ENSG00000226608 untrt    |  7.53e2 | pseudogene   ENSG00… FTLP3          540
  …        …          …          …           …           …      …            …
n-4 ENSG00000224945 trt      |  3.20e1 | sense_intro… ENSG00… RP11-…        1038
n-3 ENSG00000269958 trt      |  3.35e1 | sense_intro… ENSG00… RP11-…         811
n-2 ENSG00000260336 trt      |  5.53e1 | sense_overl… ENSG00… RP11-…        3921
n-1 ENSG00000261625 trt      |  2.21e1 | sense_overl… ENSG00… RP11-…        1284
n   ENSG00000261597 trt      |  5.01e0 | sense_overl… ENSG00… RP11-…        4094
#   n = 52
#   8 more variables: baseMean <dbl>, dispersion <dbl>, dex_trt_vs_untrt <dbl>,
#   LRTPvalue <dbl>, padj <dbl>, lfc_sign <dbl>, `` <>, dex <fct>
```

The above approach may not always be ideal if the order of the summarized columns is not known in advance. However, because we had renamed the column dimension with `cols(.samples = unique(dex))`, we may use the column names as a specific index. For example, you could use the `.assays_asis` variant as shown below.

```r
de_tpm_summary_slice <- de_tpm_summary |>
  group_by(rows(gene_biotype)) |>
  arrange(rows(
    {
      mat <- .assays_asis$ave_TPM
      abs(mat[, "trt"] - mat[, "untrt"])
    } |>
     dplyr::desc()
  )) |>
  slice(rows(1:3))

de_tpm_summary_slice
```

```
# A SummarizedExperiment-tibble Abstraction: 26 × 2
# Groups: rows(gene_biotype)
    .features       .samples | ave_TPM | gene_biotype gene_id symbol gene_length
    <chr>           <chr>    |   <dbl> | <chr>        <chr>   <chr>        <int>
  1 ENSG00000087086 untrt    |  4.47e4 | protein_cod… ENSG00… FTL            878
  2 ENSG00000075624 untrt    |  2.76e3 | protein_cod… ENSG00… ACTB          3256
  3 ENSG00000103888 untrt    |  1.79e3 | protein_cod… ENSG00… KIAA1…        7621
  4 ENSG00000269378 untrt    |  2.26e3 | pseudogene   ENSG00… ITGB1…         609
  5 ENSG00000226608 untrt    |  7.53e2 | pseudogene   ENSG00… FTLP3          540
  …        …           …             …          …          …       …           …
n-4 ENSG00000224945 trt      |  3.20e1 | sense_intro… ENSG00… RP11-…        1038
n-3 ENSG00000269958 trt      |  3.35e1 | sense_intro… ENSG00… RP11-…         811
n-2 ENSG00000260336 trt      |  5.53e1 | sense_overl… ENSG00… RP11-…        3921
n-1 ENSG00000261625 trt      |  2.21e1 | sense_overl… ENSG00… RP11-…        1284
n   ENSG00000261597 trt      |  5.01e0 | sense_overl… ENSG00… RP11-…        4094
#   n = 52
#   8 more variables: baseMean <dbl>, dispersion <dbl>, dex_trt_vs_untrt <dbl>,
#   LRTPvalue <dbl>, padj <dbl>, lfc_sign <dbl>, `` <>, dex <fct>
```

For completeness, we note that this difference could also be computed on the reshaped data, `map_dbl(.assays$ave_TPM, ~ .x["trt"] - .x["untrt"])`, though the matrix subtraction above is expected to be faster.

We can make a plot of top 3 TPM differences per biotype, with a point for the mean and a line representing the range within the group:

```r
de_tpm |>
  filter(rows(gene_biotype %in% biotypes)) |>
  filter(rows(gene_id %in% rownames(de_tpm_summary_slice))) |>
  se() |>
  ggplot(aes(x = TPM + 0.1, y = symbol, group = dex, color = dex)) +
  stat_summary(
    fun.min = min, fun.max = max, fun = mean,
    geom = "pointrange", position = position_dodge(width = 1)
  ) +
```

```r
  facet_grid(rows = vars(gene_biotype), scales = "free_y") +
  scale_x_log10(labels = scales::label_log())
```

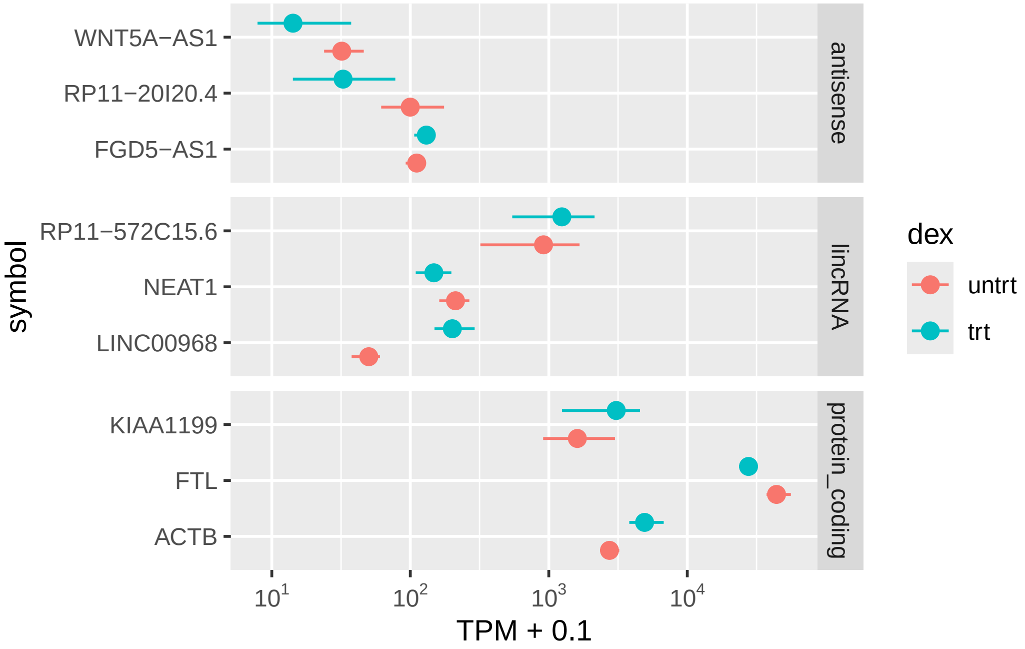


We can make a simple heatmap, additionally showing the value per donor cell line.

```r
de_tpm |>
  filter(rows(gene_biotype %in% biotypes)) |>
  filter(rows(gene_id %in% rownames(de_tpm_summary_slice))) |>
  se() |>
  ggplot(aes(x = dex, y = symbol)) +
  geom_tile(aes(fill = log10(TPM + 0.1))) +
  facet_grid(rows = vars(gene_biotype), cols = vars(cell), scales = "free_y") +
  scale_fill_viridis_c()
```

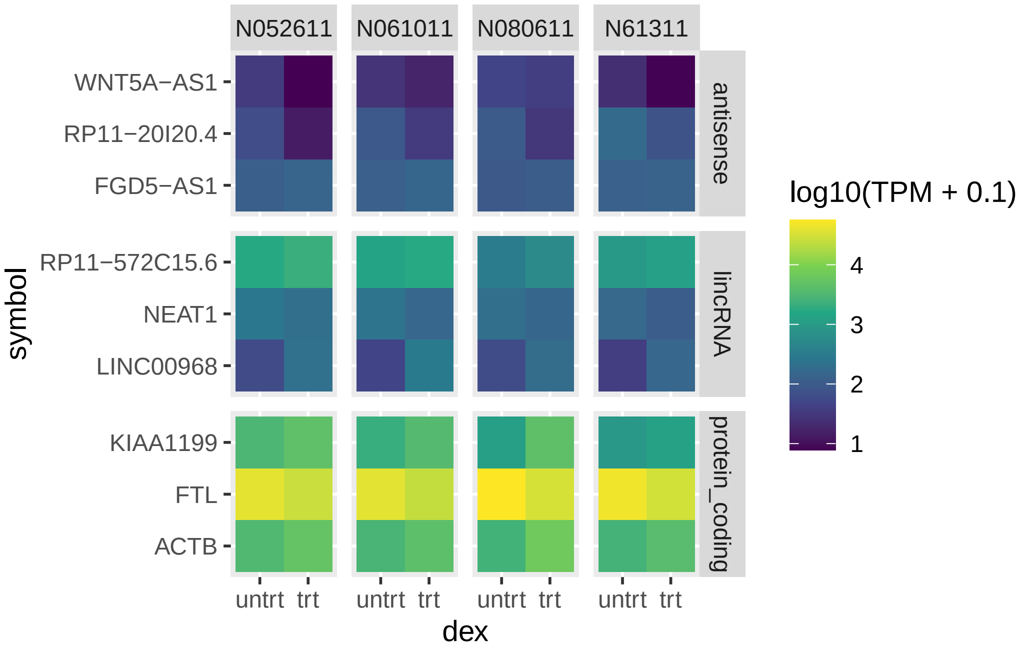


### 7 Session Info

```r
sessionInfo()
```

```
R version 4.6.0 (2026-04-24)
Platform: aarch64-apple-darwin23
Running under: macOS Sequoia 15.6.1

Matrix products: default
BLAS:   /Library/Frameworks/R.framework/Versions/4.6/Resources/lib/libRblas.0.dylib
LAPACK: /Library/Frameworks/R.framework/Versions/4.6/Resources/lib/libRlapack.dylib;  LAPACK version 3.[illegible]

locale:
[1] en_US.UTF-8/en_US.UTF-8/en_US.UTF-8/C/en_US.UTF-8/en_US.UTF-8

time zone: America/New_York
tzcode source: internal

attached base packages:
[1] stats4    stats     graphics  grDevices utils     datasets  methods
[8] base

other attached packages:
 [1] ggplot2_4.0.3                tidyr_1.3.2
 [3] dplyr_1.2.1                  tidySummarizedExperiment_1.22.0
 [5] ttservice_0.5.3              DESeq2_1.52.0
 [7] tibble_3.3.1                 plyxp_1.6.1
 [9] airway_1.32.0                SummarizedExperiment_1.42.0
[11] Biobase_2.72.0               GenomicRanges_1.64.0
```

```
[13] Seqinfo_1.2.0                 IRanges_2.46.0
[15] S4Vectors_0.50.0              BiocGenerics_0.58.0
[17] generics_0.1.4                MatrixGenerics_1.24.0
[19] matrixStats_1.5.0

loaded via a namespace (and not attached):
 [1] gtable_0.3.6        xfun_0.57           htmlwidgets_1.6.4
 [4] lattice_0.22-9      vctrs_0.7.3         tools_4.6.0
 [7] parallel_4.6.0      pkgconfig_2.0.3     Matrix_1.7-5
[10] data.table_1.18.4   RColorBrewer_1.1-3  S7_0.2.2
[13] lifecycle_1.0.5     stringr_1.6.0       compiler_4.6.0
[16] farver_2.1.2        tinytex_0.59        codetools_0.2-20
[19] htmltools_0.5.9     yaml_2.3.12         lazyeval_0.2.3
[22] plotly_4.12.0       pillar_1.11.1       ellipsis_0.3.3
[25] BiocParallel_1.46.0 DelayedArray_0.38.1 abind_1.4-8
[28] tidyselect_1.2.1    locfit_1.5-9.12     digest_0.6.39
[31] stringi_1.8.7       purrr_1.2.2         labeling_0.4.3
[34] fastmap_1.2.0       grid_4.6.0          cli_3.6.6
[37] SparseArray_1.12.2  magrittr_2.0.5      S4Arrays_1.12.0
[40] utf8_1.2.6          withr_3.0.2         scales_1.4.0
[43] rmarkdown_2.31      XVector_0.52.0      httr_1.4.8
[46] otel_0.2.0          evaluate_1.0.5      knitr_1.51
[49] viridisLite_0.4.3   rlang_1.2.0         Rcpp_1.1.1-1.1
[52] glue_1.8.1          jsonlite_2.0.0      R6_2.6.1
```
